# Two groups of *Arabidopsis* receptor kinases preferentially regulate the growth of intraspecies pollen tubes in the female reproductive tract

**DOI:** 10.1101/2020.03.17.995555

**Authors:** Hyun Kyung Lee, Daphne R. Goring

## Abstract

In flowering plants, continuous cell-cell communication between the compatible male pollen grain/growing pollen tube and the female pistil is required for successful sexual reproduction. In *Arabidopsis thaliana*, the later stages of this dialogue are mediated by several peptide ligands and receptor kinases that guide pollen tubes to the ovules for the release of sperm cells. Despite a detailed understanding of these processes, a key gap remains on the nature of the regulators that function at the earlier stages. Here, we report on two groups of *A. thaliana* receptor kinases, the *LRR-VIII-2 RK* subclass and the *SERK*s, that function in the female reproductive tract to regulate the compatible pollen grains and early pollen tube growth, both essential steps for the downstream processes leading to fertilization. Multiple *A. thaliana LRR-VIII-2 RK* and *SERK* knockout mutant combinations were created, and several phenotypes were observed such as reduced wild-type pollen hydration and reduced pollen tube travel distances. As these mutant pistils displayed a wild-type morphology, the observed altered responses of the wild-type pollen are proposed to result from the loss of these receptor kinases leading to an impaired pollen-pistil dialogue at these early stages. Furthermore, using pollen from related Brassicaceae species, we also discovered that these receptor kinases are required in the female reproductive tract to establish a reproductive barrier to interspecies pollen. Thus, we propose that the *LRR-VIII-2 RK*s and the *SERK*s play a dual role in the preferential selection and promotion of intraspecies pollen over interspecies pollen.

## Introduction

In flowering plants, cell-cell communication is a required component of coordinating the directional growth of pollen tubes which are delivering sperm cells to the female gamete-containing ovules deeply embedded in the pistil [1-5]. The female reproductive tract in the pistil can also act as a selective ‘sieve’ to impose physical barriers to non-compatible pollen tubes, for example, from other species [6-9]. In *A. thaliana*, these regulatory constraints are rapidly imposed following pollination as the desiccated pollen grain is reliant on the stigma, at the top of the pistil, to release water and other components required for hydration, germination and pollen tube entry into the stigma barrier [1, 8, 10]. Following this, the compatible pollen tube follows a long path of growing intercellularly through the densely packed tissue of the female reproductive tract, consists of the transmitting tissue of the stigma and style to the transmitting tract of the ovary, where it then emerges to find an ovule [1, 11]. Thus, proper dialogue between the pollen tube and the pistil is required for precise pollen tube navigation that would eventually lead to characteristic double fertilization.

A number of key studies have identified the *A. thaliana* peptide ligands and receptor kinases (RKs) that regulate pollen tube guidance towards the ovule and pollen tube reception at the ovule. For example, the POLLEN-SPECIFIC RECEPTOR-LIKE KINASEs (PRKs) and MALE DISCOVERER1 (MDIS1)-MDIS1-INTERACTING RECEPTOR LIKE KINASEs (MIKs) are used by the pollen tube to perceive ovular-borne AtLURE guidance peptides which orient the pollen tube towards a viable ovule [12-15]. At the ovule, pollen tube reception is mediated by a co-receptor complex, FERONIA (FER) and glycosylphosphatidylinositol-anchored protein (GPI-AP), LORELEI, located in the filiform apparatus to correctly initiate sperm cells discharge for double fertilization [16-19]. Furthermore, pollen tube integrity during its rapid polarized cell growth phase is maintained via autocrine signaling as the pollen tube uses several CrRLK1L-group receptor kinases and GPI-APs to strengthen the cell wall, thereby mitigating premature pollen tube bursting [20-22]. Thus, a detailed picture has emerged of the complex regulatory processes regulating these later stages of pollen-pistil interactions. However, what is still missing is the identification of the key regulators that function in the earlier stages that take place in the stigma/style immediately after pollination. Successful growth of pollen tubes through the *A. thaliana* stigma/style is a prerequisite to these later stages as pollen tubes growing through the stigma/style then acquire the competency to perceive ovular guidance cues [23]. So far, the only putative early stage regulators that have been identified are the pollen-specific genes encoding the *Pollen Coat Protein-B* (*PCP-B*) peptides which are required for pollen adhesion and hydration [24].

Here, we report that members of two groups of *RK* genes, the *LEUCINE RICH REPEAT (LRR)-VIII-2 RK*s and the *SOMATIC EMBRYOGENESIS RECEPTOR KINASES* (*SERKs*), that are required in the stigma and the female reproductive tract to promote compatible pollen grain hydration and pollen tube growth and to establish a reproductive barrier against interspecies pollen tube growth. The discovery of these RKs was enabled by employing an unconventional approach of using an interspecies tissue-specific pseudokinase as a tool for discovery of candidate RKs by using the assumption that pseudokinases typically associate with active kinases in a complex [25-28]. In conclusion, our findings have uncovered several RKs as regulators of the early stages of *A. thaliana* pollen-pistil interactions and provide an exciting platform for identifying additional signaling players that regulate the complex molecular dialogue leading to successful plant sexual reproduction.

## Results

### *At-RKF1, At-SERK1* and *At-SERK3/BAK1* are the interactors of a stigma-specific pseudokinase

Previously, a search for stigma-specific candidate genes failed to identify any RK genes [29, 30], and a survey of a developmental RNA-Seq transcriptome [31] for predicted *A. thaliana* RK genes confirmed the absence of any stigma-specific RK genes, other than the pseudogenized *SRKA* gene (Table S1). As many RK genes are expressed in the stigma (Table S1), we took the approach of using the stigma-specific *Arabidopsis lyrata* BRASSIKIN1 (Al-BKN1) pseudokinase [29] as a tool to narrow down the list of potential receptor kinase candidates to test for functions in the stigma following pollination. Al-BKN1 and three other BKNs were used in a yeast two-hybrid pairwise interaction screen with cytosolic kinase domains from 48 Leucine Rich Repeat (LRR)-RKs and 6 other RKs (Figure S1A and Table S2). The positive hits identified were narrowed down to two different groups of LRR-RKs (Figure 1A and S1B-E). These RKs belong to the LRR-VIII-2 RKs (At-RKF1, At-RKFL1, At-RKFL2) which have a distinct extracellular domain with LRR repeats followed by a malectin domain [32] and the LRR-II RKs (At-SERK1, At-SERK3/BAK1) which have smaller extracellular domains with LRR repeats [33]. The function of At-RKF1 [34] and its closely related paralogs (At-RKFL1, 2, 3) are unknown; however, the At-SERKs are a well-established group of co-receptors for multiple RK partners [33]. Furthermore, we observed protein interactions with RKF1 or RKFL1 homologs from *Arabidopsis lyrata, Eutrema salsugineum* and *Brassica rapa*, suggesting a conserved interaction across Brassicaceae species (Figure 1A, S1B and S1D). To validate the observed Y2H protein interactions, bimolecular fluorescence complementation (BiFC) assay was used to test for putative interactors by transiently expressing full-length proteins in *Nicotiana benthamiana* leaf epidermal cells. Confocal laser scanning microscopy (CLSM) images confirmed that Al-BKN1 interacted with At-SERK1, At-SERK3/BAK1, Al-RKF1, At-RKFL1 and Br-RKF1 A8, and the interactions were localized to the plasma membrane (Figure 1B and S2).

**Figure 1.**
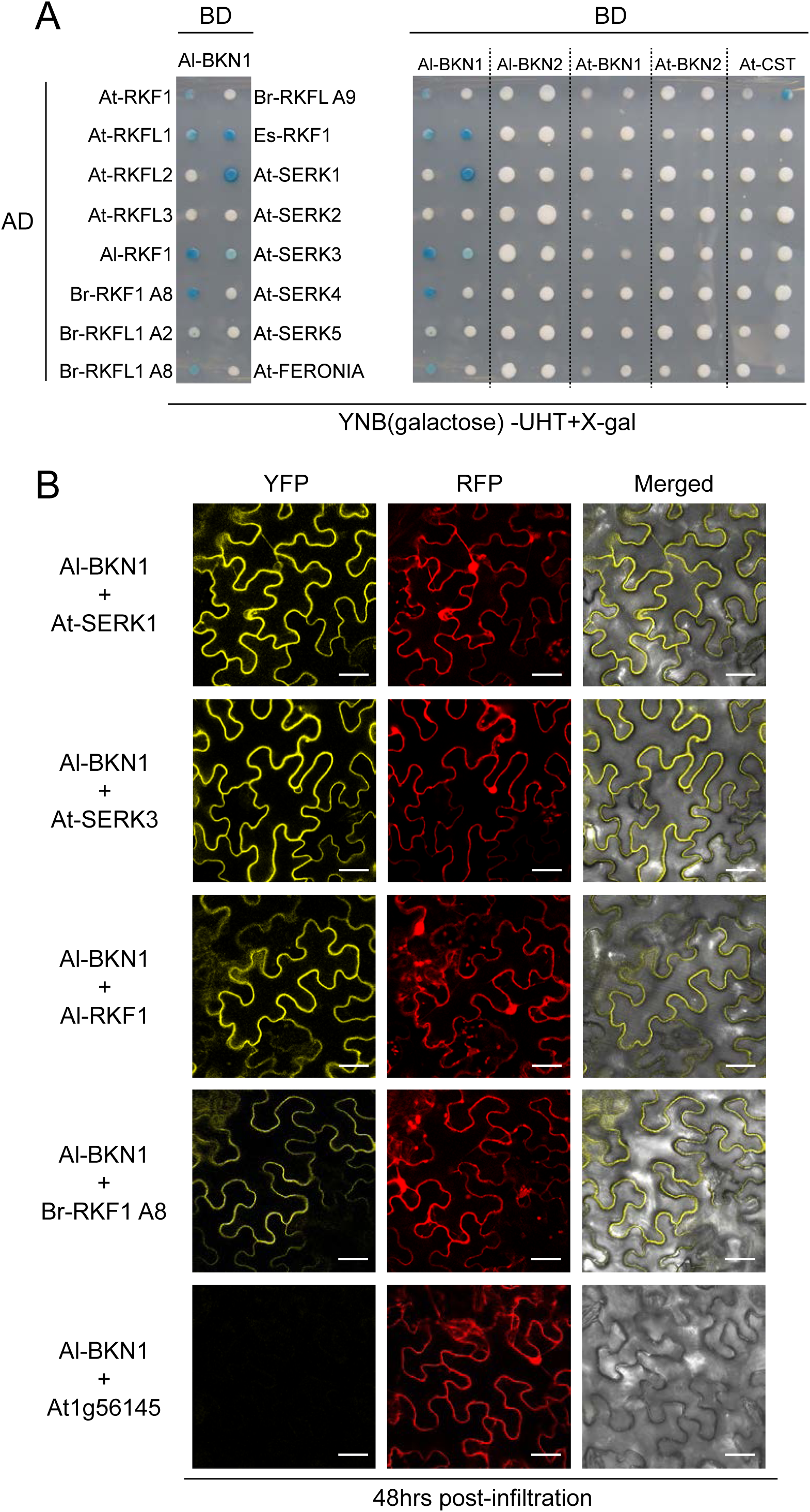
*At-RKF1, At-SERK1* and *At-SERK3/BAK1* are the interactors of a stigma-specific pseudokinase. (A) Y2H summary result for a subset of the RK cytosolic domains screened for interactions with Al-BKN1, Al-BKN2, At-BKN1 and At-BKN2. At-CST was included as a negative control. Blue coloured yeast colonies indicate positive RK interactors (At-RKF1s, At-SERKs and RKF1/RKFL1 homologs from *A. lyrata, B. rapa* or *E. salsugineum*); See also Figure. S1 for the full screen of 54 different RKs with different BKNs. (B) Confirmation of positive interactors using the BiFC assay with full-length proteins. The YFP signal indicates positive protein-protein interactions which are localized to the plasma membrane as predicted for RKs; see Figure S2 for additional images. RFP is an internal control for transformation and is localized to both the cytoplasm and nucleus. CLSM images of abaxial leaf epidermis of *N. benthamiana* were taken 48-hours after *A. tumefaciens*-mediated infiltration. Scale bar = 30 µm.

### *At-RKF1* and paralogs control wild-type pollen hydration rate

The observed protein interactions with different Brassicaceae RKF1 homologs potentially indicated a conserved novel function for these RKs in the stigma, and so this was investigated through *A. thaliana* knockout mutants. As the *At-RKF1* gene is present in a cluster with *At-RKFL1, At-RKFL2* and *At-RKFL3*, a 25 kb cluster deletion was created (Figure 2A). Two independent quadruple mutants, *rkfΔ-1* and *rkfΔ-2*, displayed an overall wild-type plant and flower morphology (Figure 2B, S3 and S4A). The *rkfΔ-1* quadruple mutant was then crossed to *serk1-1 bak1-4* double mutant to generate the *rkfΔ-1serk1-1bak1-4* sextuple mutant. The *serk1-1 bak1-4* double mutant introduced some brassinosteroid-insensitive phenotypes previously associated with the *serk* mutants [35-37], characterized by overall reduced growth, delayed flowering, and anther filaments failing to fully elongate. However, the pistil morphology appeared wild-type in these mutants with the stigma and style formed normally (Figure 2B and S4A). Pistils from the different mutant *RK* combinations were then assessed for the ability to support the very first physiological output of compatible pollen-pistil interactions. Pollen hydration is one of the earliest post-compatible pollination steps where water is released from the *A. thaliana* stigma to the desiccated pollen grain [1, 8]. Pollen hydration assays were conducted by applying wild-type pollen grains to the mutant stigmas (Figure 2C). Both wild-type Col-0 and *serk1-1 bak1-4* stigmas supported a normal rate of wild-type Col-0 pollen hydration at 10-minutes post-pollination. In contrast, Col-0 pollen grains applied to stigmas from the two independent *rkfΔ-1* and *rkfΔ-2* quadruple mutants showed a significant reduction in hydration at 10-minutes post-pollination (Figure 2C). The same level of reduced Col-0 pollen hydration was also observed on *rkfΔ-1serk1-1bak1-4* sextuple mutant stigmas. As a control, mutants for the *FER* [16, 38] were tested for Col-0 pollen hydration, and no changes were observed (Figure S5A).

**Figure 2.**
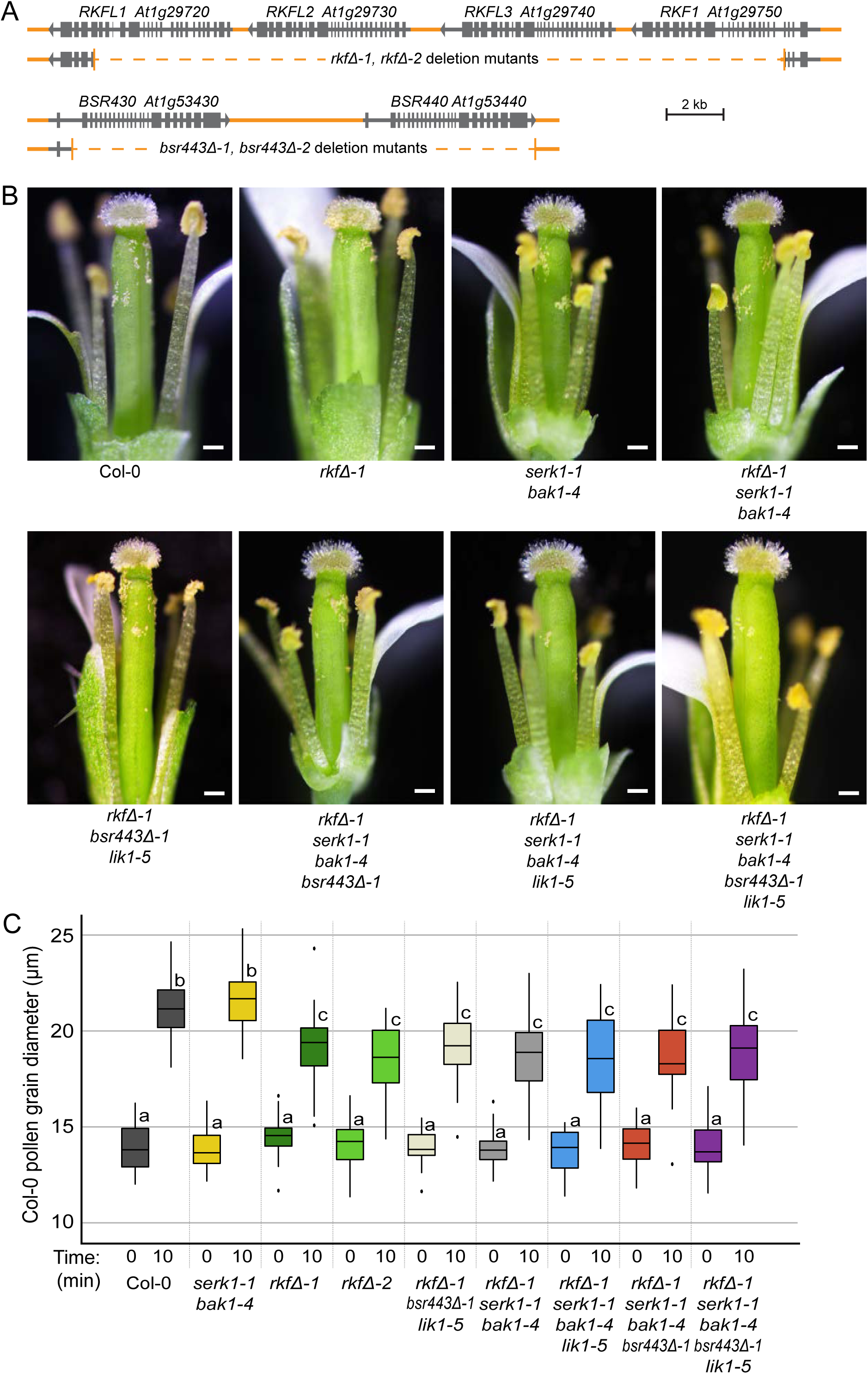
*At-RKF1* and paralogs regulate wild-type pollen hydration rate. (A) Schematic representation of CRISPR deletion mutations generated in the *At-RKF1* cluster and the tandem *At-BSR440* and *At-BSR430* genes; see Table S6 for a full list of mutants. (B) Representative images of stage-14 flowers from wild-type Col-0 and the various *RK* mutants. Some petals and sepals have been removed to better view the pistil and anthers; see Figure S4 for all mutant flower photos. Scale bar = 200 µm. (C) Pollen hydration assays for wild-type Col-0 pollen grains placed on wild-type Col-0 and mutant *RK* stigmas. Hydration was measured by taking pollen grain diameters at 0- and 10-minutes post-pollination. Wild-type levels of pollen hydration was supported by the *serk1-1bak1-4* stigmas, while there was a significant reduction at 10-minutes post-pollination in Col-0 pollen hydration on *RK* mutant stigmas that carried the *rkf* cluster deletion (*rkfΔ*). Data are plotted as box plots displaying first (Q1) and third (Q3) quartiles split by the median; the whiskers extend maximum of 1.5 times the interquartile range beyond the box. Outliers are indicated as black dots. n=30 pollen grains per line, P<0.05 (One-way ANOVA with Tukey-HSD post-hoc test).

Additional higher order *RK* mutants were then created by selecting other LRR-VIII-2 RK members that displayed high expression in the stigma: *LIK1/BSR840, BSR440* and *BSR430/NILR2* [39-41] (Figure S1E and Table S1-S3). The *BSR440* and *BSR430* genes are tandemly linked and so both genes were deleted using CRISPR to generate the *bsr443Δ* double mutant (Figure 2A). The *rkfΔ-1serk1-1bak1-4lik1-5* septuple mutant, *rkfΔ-1serk1-1bak1-4bsr443Δ-1* octuple mutant, and the *rkfΔ-1serk1-1bak1-4bsr443Δ-1lik1-5* nonuple mutant were similar in overall appearance and only displayed the *serk1-1bak1-4* associated-mutant traits (Figure 2B, S3 and S4). In contrast, the *rkfΔ-1lik1-5bsr443Δ-1* septuple mutant (*LRR-VIII-2* only mutant alleles) displayed a wild-type floral and overall plant morphology. Importantly, none of the higher order mutants showed developmental or morphological defects at the level of stigma and style that could potentially impact compatible pollen interactions (Figure 2B). Stigmas from all these higher order mutant combinations were tested using the pollen hydration assay and were found to support a similar level of reduced wild-type Col-0 pollen hydration rates as seen for the *rkfΔ* quadruple mutants (Figure 2C). This indicated that among these RKs, the *At-RKF1* gene cluster is primarily responsible for regulating compatible pollen hydration by the stigma.

### Higher order mutant pistils cause altered wild-type pollen tube growth patterns

Despite the reduced pollen hydration rate at 10-minutes post-pollination on the *rkfΔ* quadruple mutant stigmas, the Col-0 pollen grains were observed to germinate and produce pollen tubes. To assess if this next stage of the compatible pollen-pistil interactions was impacted in the *RK* mutant pistils, wild-type pollen tube growth through the mutant stigma/styles was observed. At 2-hours post-pollination, pistils were fixed and stained with the aniline blue that stains for callose and allows for the visualization of pollen tubes (Figure 3A-3H and S6). Wild-type Col-0 pollen tubes were observed to have grown through the stigma/style and into the transmitting tract of pistils from Col-0, the lower level *RK* mutant combinations (*rkfΔ-1* quadruple mutant and *serk1-1bak1-4* double mutant), and the *rkfΔ-1lik1-5bsr443Δ-1* septuple mutant (Figure 3A-3D and S6).

**Figure 3.**
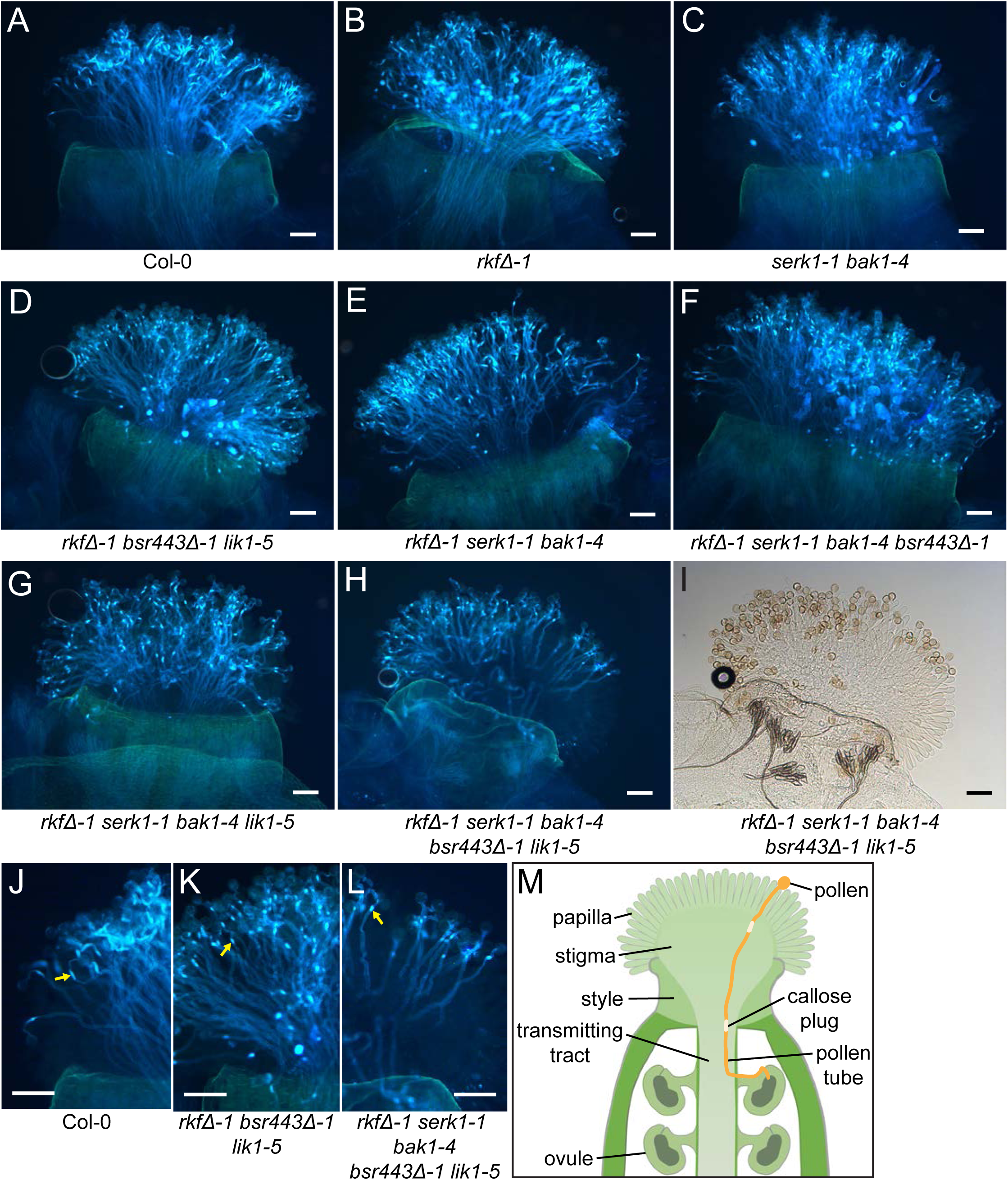
Higher order mutant pistils cause altered wild-type pollen tube growth patterns. (A-H) Representative images of pollinated pistils stained with aniline blue. Col-0 pollen grains were applied to stigmas from Col-0 or *RK* mutants and the pollinated pistils were fixed 2-hours post-pollinations. The images show the stigmas, styles and upper part of the transmitting tract of the pollinated pistils. Wild-type pollen tubes grew a shorter distance on the higher order mutant stigmas. Number of pistils pollinated for each line: n=10 and replicates show similar results; scale bar = 100 µm. (I) Brightfield image for (H) showing the stigma and pollen grains; see Figure S6 for the other corresponding brightfield images. (J-L) Close-up images of aniline blue stained pistils to better show the different morphologies of the callose plugs formed in Col-0 pollen tubes grown on Col-0 stigmas versus the higher order mutants (see yellow arrows). Scale bar = 100 µm. (M) Schematic of the top part of an *A. thaliana* pistil with the different parts labelled.

Interestingly, wild-type Col-0 pollen tubes on the higher order *RK* mutant pistils grew shorter distances when *serk1-1bak1-4* was combined with the different *LRR-VIII-2 RK* mutants (*rkfΔ-1serk1-1bak1-4* sextuple mutant, *rkfΔ-1serk1-1bak1-4lik1-5* septuple mutant, *rkfΔ-1serk1-1bak1-4bsr443Δ-1* octuple mutant, *rkfΔ-1serk1-1bak1-4bsr443Δ-1lik1-5* nonuple mutant). This was most noticeable with the *rkfΔ-1serk1-1bak1-4bsr443Δ-1lik1-5* nonuple mutant pistils, where wild-type Col-0 pollen tubes had only reached the style at 2-hours post-pollination (Figure 3H and S6). Furthermore, higher order *RK* mutant pistils show differences in the morphology of callose plugs within the wild-type Col-0 pollen tubes (Figure 3K-3L). Callose plugs are deposited to separate the growing pollen tube tip from the older part of the tube, and they are typically elongated in shape at 2-hours post-pollinations [42, 43] as seen for the Col-0 pollen tubes growing in Col-0 pistils and some of the *RK* mutant pistils (Figure 3A-3C, 3E and 3J). However, in some of the higher order mutants, the callose plugs were not elongated (Figure 3D, 3F-3H, 3K and 3L). This did not appear to be linked to reduced pollen tube growth as the wild-type Col-0 pollen tubes also displayed this smaller callose plug phenotype on the *rkfΔ-1lik1-5bsr443Δ-1* septuple mutant pistils (Figure 3D and 3K). Thus, the formation of normal callose plugs in wild-type pollen tubes is linked to the action of these LRR-VIII-2 RKs in the stigma and style.

### *At-SERK1, At-SERK3/BAK1* and *LRR-VIII-2 RKs* are required for efficient pollen tube growth and seed production

To further investigate the observation that wild-type Col-0 pollen tubes were travelling shorter distances in the higher-order mutant pistils, Col-0 *Lat52p:GUS* pollen grains were used to allow for GUS staining and quantification [44]. Pistils from Col-0 and the different *RK* mutant combinations were pollinated with the Col-0 *Lat52p:GUS* pollen and left for 2- or 6-hours (Figure 4A, 4B and S7). Consistent with the aniline blue results, the pollen tubes traveled well into the transmitting tract at 2-hours post-pollination in Col-0 and the *rkfΔ-1* and *rkfΔ-2* quadruple mutant pistils, with a number of ovules being reached by 6-hours post-pollination (Figure 4A, 4B and S7). For the *serk1-1bak1-4* double mutant pistils and the *rkfΔ-1lik1-5bsr443Δ-1* septuple mutant pistils, significant reductions in the Col-0 *Lat52p:GUS* pollen tube travel distances were observed at 6-hours post-pollination (Figure 4B). Further significant reductions in pollen tube growth distances were observed when different *LRR-VIII-2 RK* mutants were combined with *serk1-1bak1-4* indicating a synergistic effect of these SERKs and LRR-VIII-2 RKs on pollen tube growth in the upper pistil (Figure 4B). The strongest decrease in Col-0 *Lat52p:GUS* pollen tube growth distances was seen in the *rkfΔ-1serk1-1bak1-4bsr443Δ-1lik1-5* nonuple mutant pistils where the pollen tubes traveled significantly shorter distances at both 2- and 6-hours post-pollination (Figure 4A and 4B). Thus, wild-type pollen tubes require the action of these SERK and LRR-VIII-2 RKs in the transmitting tissue of the upper pistil to grow efficiently.

**Figure 4.**
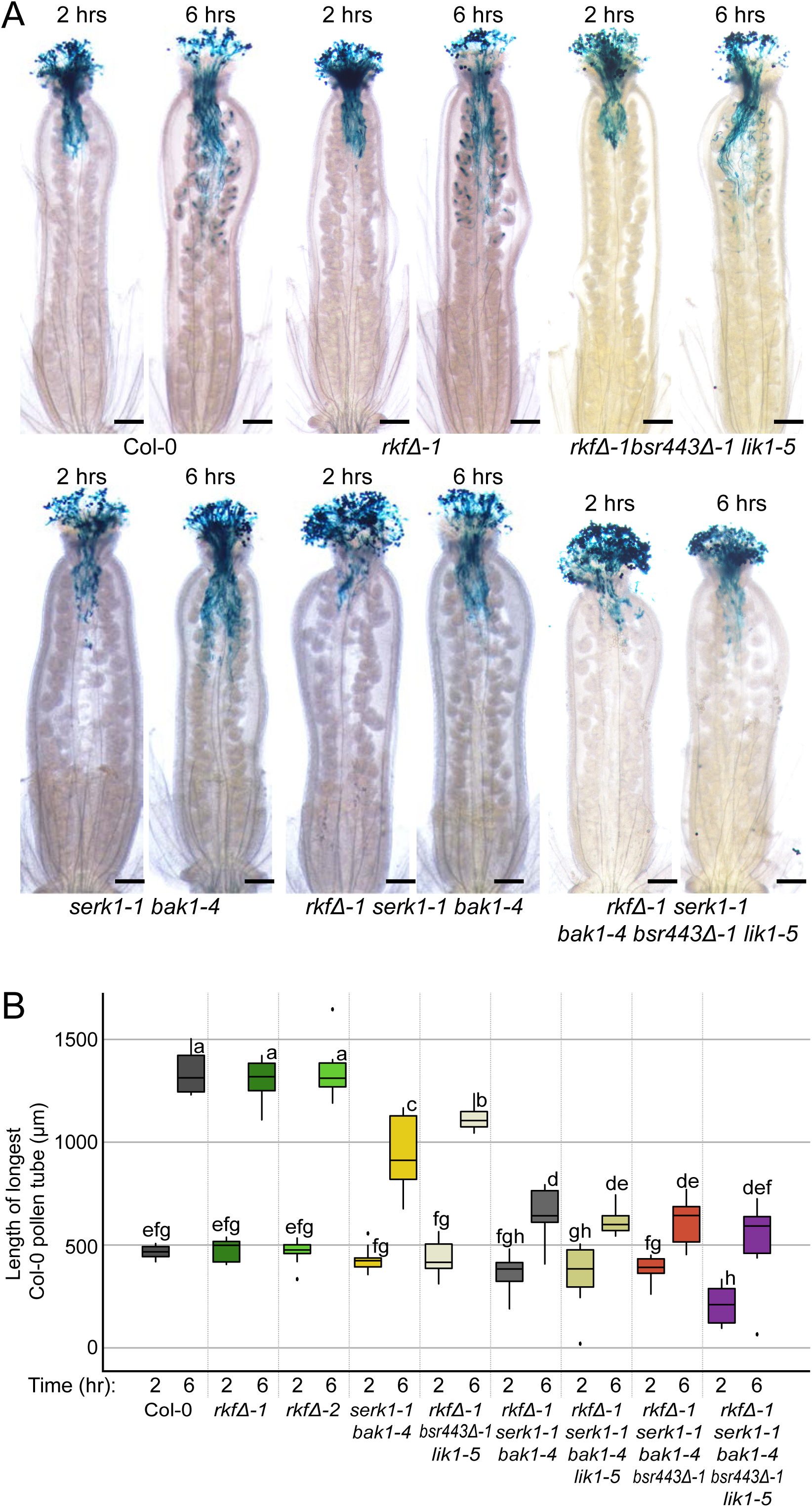
*At-SERK1, At-SERK3/BAK1* and *LRR-VIII-2 RKs* are required for efficient pollen tube growth. (A) Representative images of pistils pollinated with Col-0 *Lat52p:GUS* pollen [44] for 2- or 6-hours and then fixed and stained for GUS activity; see also Figure S7 for additional images. Scale bar = 200 µm. (B) Distance travelled by Col-0 *Lat52p:GUS* pollen tubes in the Col-0 and various *RK* mutant pistils. Pollen tube growth was measured by drawing a line across the bottom of the style and measuring the distance between that line to the leading GUS-stained pollen tube tip [44]. All *RK* mutants except *rkfΔ-1* and *rkfΔ-2* lines showed shorter travel distances at 6-hours post-pollinations. The nonuple mutant pistils also displayed a reduction at 2-hours post-pollinations. Data are plotted as box plots displaying first (Q1) and third (Q3) quartiles split by the median; the whiskers extend maximum of 1.5 times the interquartile range beyond the box. Outliers are indicated as black dots. n = 10 pistils per time point, P<0.05 (One-way ANOVA with Tukey-HSD post-hoc test).

Pollen tube growth is an essential step before the later stages of delivering sperm cells to an ovule for fertilization, and as a result, the effect of knocking these *SERK* and *LRR-VIII-2 RK* genes in the pistil could also impact seed set. To address this, pistils from Col-0 and the various *serk1-1bak1-4 LRR-VIII-2 RK* mutant combinations were manually pollinated with wild-type Col-0 pollen grains and left for 2-weeks prior to counting seeds. Siliques from the *rkfΔ-1* and *rkfΔ-2* quadruple mutants, along with the *rkfΔ-1lik1-5bsr443Δ-1* septuple mutant, produced similar seed counts compared to Col-0. However, significant reductions in seed counts were measured for the remaining *RK* mutant combinations (Figure 5A). In the case of the *serk1-1bak1-4* double mutant, this could be attributed to a reduction in the numbers of ovules in the mutant pistils where equivalent values of ∼46 ovules/pistil and ∼46 seeds/silique were measured (Figure 5A and 5B). The number of ovules per pistil remained the same when the *LRR-VIII-2 RK* mutant combinations were combined with *serk1-1bak1-4*, but importantly, the number of seeds per silique were further reduced for these higher order mutants (Figure 5A and 5B). Again, the *rkfΔ-1serk1-1bak1-4bsr443Δ-1lik1-5* nonuple mutant had the strongest phenotype, exhibiting the lowest average seed set (∼28 seeds/silique). A closer inspection of the siliques indicated that unfertilized ovules contributed to the reduction in seed counts suggesting that pollen tubes had failed to reach these ovules for fertilization (Figure 5C).

**Figure 5.**
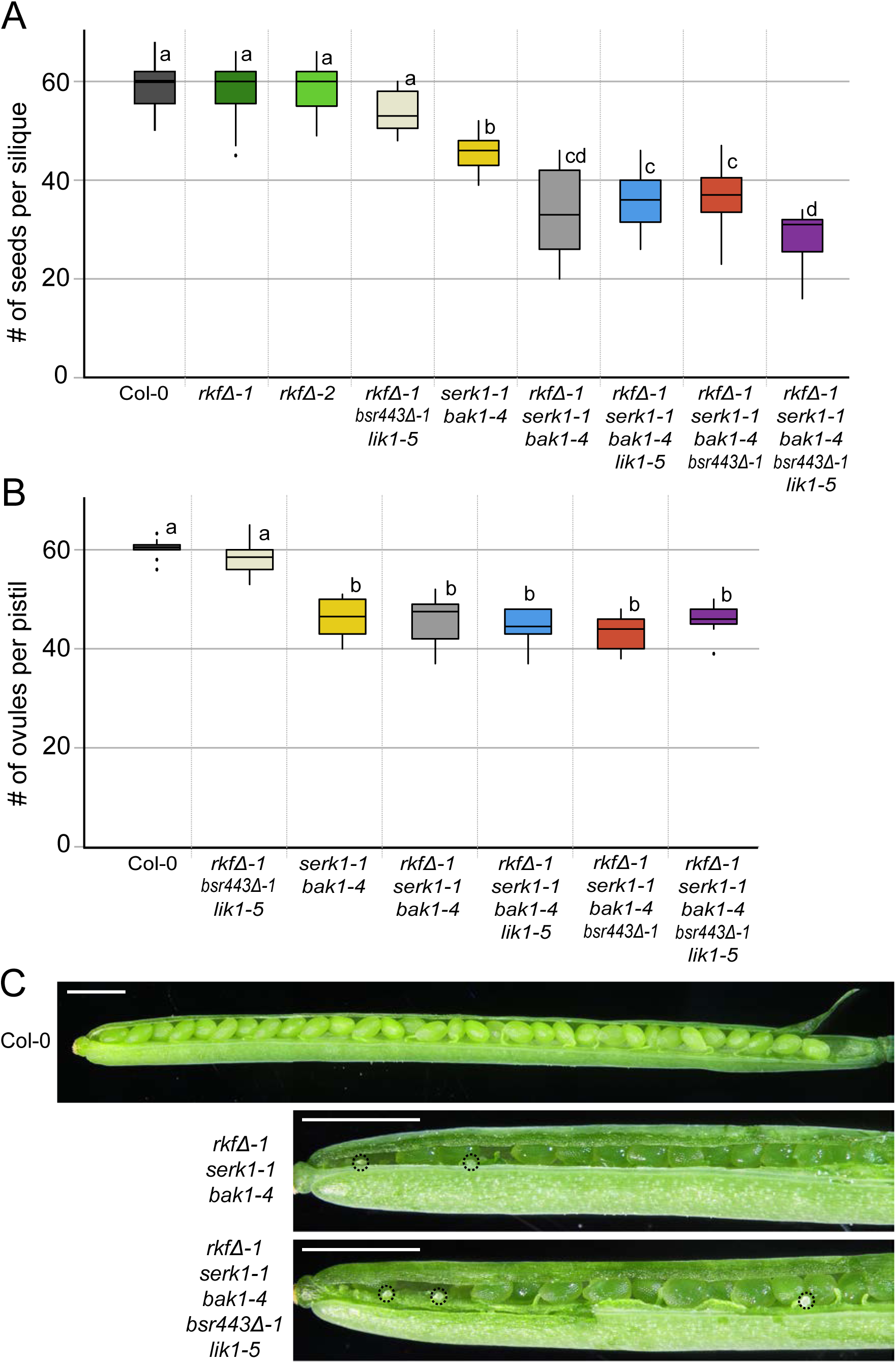
Seed and ovule counts for the different *RK* mutant combinations. (A) Seeds per silique counts following manual pollination with wild-type Col-0 pollen. Pistils from Col-0 or the *RK* mutants were manually pollinated with Col-0 pollen and seeds per siliques were counted at 2-weeks post-pollinations. Significant reduction in the number of seeds per silique was observed for the *RK* mutant pistils, except for *rkfΔ-1, rkfΔ-2* and *rkfΔ-1bsr443Δ-1lik1-5*. n = 15 siliques per line, P<0.05 (One-way ANOVA with Tukey-HSD post-hoc test). (B) Ovule numbers per pistil in Col-0 and the *RK* mutants. All pistils that contained *serk1-1bak1-4* show reduced number of ovules per pistil. n = 10 pistils per line, P<0.05 (One-way ANOVA with Tukey-HSD post-hoc test). (A and B) Data are plotted as box plots displaying first (Q1) and third (Q3) quartiles split by the median; the whiskers extend maximum of 1.5 times the interquartile range beyond the box. Outliers are indicated as black dots. (C) Representative images of manually pollinated siliques for Col-0, *rkfΔ-1serk1-1bak1-4*, and *rkfΔ-1serk1-1bak1-4bsr443Δ-1lik1-5*. Siliques were cracked open for further observations, and dotted circles indicate unfertilized ovules. Replicates showed similar results. n = 5 siliques per line. Scale bar = 1.0 mm.

### Interspecies pollinations reveal increased pollen tube entry into higher order mutant pistils

Signaling proteins required for pollen-pistil interactions can also have additional roles in promoting self-pollen over pollen from related species [9, 15]. To investigate if the *SERKs* and *LRR-VIII-2 RKs* could also have this function in the pistil, we conducted interspecies pollinations using two Brassicaceae species, the closely-related *C. rubella* and the more distantly-related *E. salsugineum*. Both *C. rubella* and *E. salsugineum* self-pollinations are fully compatible with lots of pollen tube growth (Figure 6A and 6E). In contrast, when *C. rubella* pollen were applied to Col-0 pistils, very few germinated and only a few pollen tubes were seen in the style with an average compatibility score of 2.1 as previously reported [9] (Figure 6B and 6I). *E. salsugineum* pollen applied to Col-0 pistils similarly resulted in very low numbers of pollen tubes in the style with a compatibility score of 1.9, but most pollen grains germinated on the papillae surface forming short pollen tubes (Figure 6F and 6I). Interestingly, the *rkfΔ-1bsr443Δ-1lik1-5* septuple mutant pistils supported many more *C. rubella* and *E. salsugineum* pollen tubes growing into the style with a significant increase in the compatibility scores to 3.6 and 3.3, respectively (Figure 6C, 6G and 6I). The *rkfΔ-1serk1-1bak1-4bsr443Δ-1lik1-5* nonuple mutant pistils displayed mixed results with a very low number of *C. rubella* pollen tubes in the style similar to Col-0 (compatibility score of 2.0, Figure 6D, 6I), while a significant increase in *E. salsugineum* pollen tubes were again observed similar to *rkfΔ-1bsr443Δ-1lik1-5* (compatibility score of 3.8, Figure 6G, 6I). It was unclear why the addition of the *serk1-1bak1-4* alleles to the *LRR-VIII-2 RKs* mutant combinations somehow reversed this effect for *C. rubella* pollinations. Overall, this suggests that the *LRR-VIII-2 RKs* have a dual function in promoting intraspecies pollen and presenting a barrier to interspecies pollen.

**Figure 6.**
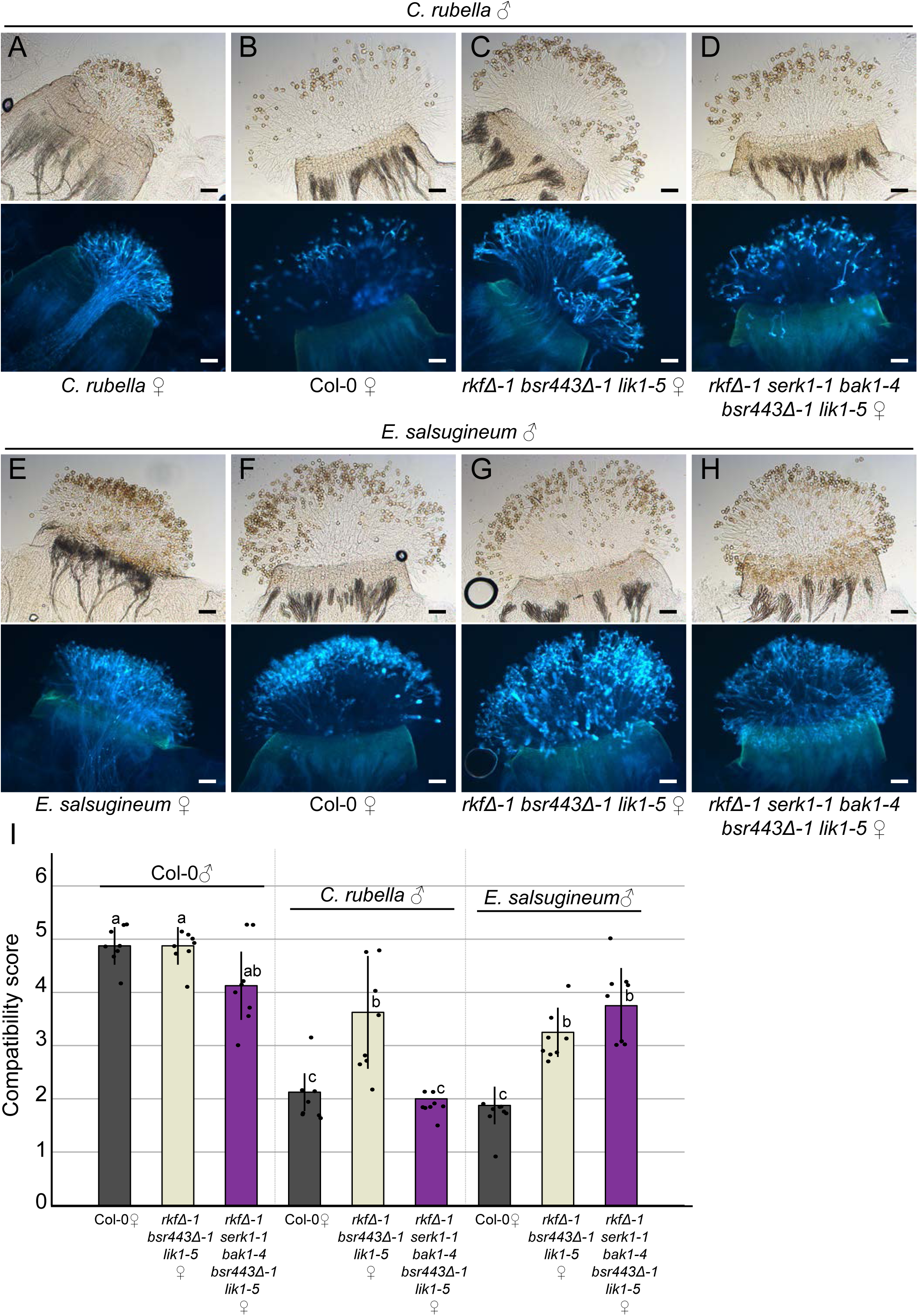
Interspecies pollinations reveal increased pollen tube entry into higher order mutant pistils. (A-D) Representative images of brightfield (top) and aniline blue stained (bottom) pistils from *C. rubella*, Col-0, *rkfΔ-1bsr443Δ-1lik1-5*, and *rkfΔ-1serk1-1bak1-4bsr443Δ-1lik1-5* manually pollinated with *C. rubella* pollen. (E-H) Representative images of brightfield (top) and aniline blue stained (bottom) pistils from *E. salsugineum*, Col-0, *rkfΔ-1bsr443Δ-1lik1-5*, and *rkfΔ-1serk1-1bak1-4bsr443Δ-1lik1-5* manually pollinated with *E. salsugineum* pollen. All pistils were fixed at 6-hours post-pollinations, and replicates showed similar results. n = 8 pistils per line. Scale bar = 100 µm. (I) Compatibility scores of Col-0, *rkfΔ-1bsr443Δ-1lik1-5*, and *rkfΔ-1serk1-1bak1-4bsr443Δ-1lik1-5* pistils pollinated with Col-0, *C. rubella* or *E. salsugineum* pollen. The scores were measured as previously described [9]. Black dots represents a single data point; the plot has been jittered to visualize all data points. Data are represented as mean ± SD; bar represent mean values and whiskers represent SD. n = 8 pistils per line, P<0.05 (One-way ANOVA with Tukey-HSD post-hoc test).

## Discussion

Previous work in the field has uncovered several key RKs and peptide ligands that regulate *A. thaliana* reproduction; however, these players function at later stages such as in ovular pollen tube guidance and pollen tube reception [1-5]. Little is known about the identity of signaling components in the upper regions of the pistil regulating pollen-pistil interactions. Here, we have discovered two groups of *A. thaliana RK*s, the *SERK*s and *LRR-VIII-2 RK*s, that function together in the stigma and style to regulate wild-type pollen tube growth. Our data indicates that the *At-RKF1* cluster is required in the stigma for full hydration of wild-type pollen. Moreover, combinations of *RK* mutants up to the *rkfΔ-1serk1-1bak1-4bsr443Δ-1lik1-5* nonuple mutant implicate both the *SERK*s and *LRR-VIII-2 RK*s acting in the upper pistil to support wild-type pollen tube growth and proper callose plug formation. These two groups can separately function in the reproductive tract to support pollen tube growth as seen in the pollinated *serk1-1bak1-4* double mutant pistils and the *rkfΔ-1lik1-5bsr443Δ-1* septuple mutant pistils. However, further reductions in wild-type pollen tube travel distances in the stigma and style tissues only began to appear when the *rkfΔ-1* quadruple mutations were combined with *serk1-1bak1-4*, with the strongest reduction seen in the *rkfΔ-1serk1-1bak1-4bsr443Δ-1lik1-5* nonuple mutant pistils. The synergistic effects seen when the *A. thaliana SERK*s and *LRR-VIII-2 RK*s mutants were combined suggest that they act independently, which would be consistent with a previous *in vitro* interactome dataset that did not find any interactions between their extracellular domains [45]. Finally, smaller callose plugs were observed in the wild-type pollen tubes growing through the higher order *RK* mutant pistils. Pollen tubes sustain rapid polarized growth by depositing callose plugs to maintain turgor pressure and concentrate the cytoplasmic content at the growing tip [43, 46]. The loss of the signaling dialog between the pollen tube and these RKs in the stigma/style transmitting tissue may cause the wild-type pollen tube to have reduced access to components needed for proper callose plug formation. Interestingly, the altered callose plug morphology is similar in appearance to that seen in *in vitro* grown pollen tubes [47].

In addition to having a role in supporting *A. thaliana* pollen tube growth in the upper pistil, we also discovered that the *LRR-VIII-2 RK*s also function in presenting a barrier to interspecies pollen tubes [9, 15]. When pollen grains from *C. rubella* or *E. salsugineum* were applied to the *LRR-VIII-2 RK*s mutant pistils, growth of these interspecies pollen tubes through the stigma/style tissues was observed, while wild-type Col-0 pistils efficiently blocked interspecies pollen tube growth. Presenting a barrier to pollen from other species is an important trait as it allows the pistil to control mate selection and increase the rate of successful fertilization, but the impact of such as system should also be viewed in the context of mating patterns for a particular flowering species. For example, *A. thaliana* is a predominantly selfing specie and has evolved floral traits that promote self-pollen deposition on the stigma [48]. In contrast, *A. lyrata* and *A. halleri* are predominantly outcrossing species and rely on insect pollinators to deliver pollen, potentially from many different sources [48]. Thus, the *LRR-VIII-2 RK* interspecies barrier would be predicted to have a bigger impact in these outcrossing *Arabidopsis* species, and perhaps it is in this context where the observed Al-BKN1/Ah-BKN1 interactions with these RKs comes into play, a direction for future investigations.

Overall, our data has provided new insights on the roles of *A. thaliana* RKs in regulating pollen-pistil interactions with the identification of two new groups of *RK* genes that are required in the upper pistil for successful compatible pollen interactions. We have shown that the At-RKF1 and its paralogs function in the stigma to regulate pollen hydration while the addition of other *A. thaliana* LRR-VIII-2 RKs together with At-SERK1 and At-SERK3/BAK1 function in the stigma/style regions to promote pollen tube travel. Thus, these findings further highlight the importance of pistil-expressed *RK*s to mediate a dialogue between the pollen/pollen tube and the pistil. Finally, our work has simultaneously implicated the *A. thaliana* LRR-VIII-2 RKs in both promoting self-pollen tube growth and presenting a barrier for interspecific pollen from other Brassicaceae species.

## Acknowledgements

We thank Safa Abdulsalam, Paula Beronilla and Lewis Kurschner for technical assistance, and members of the Goring lab for critically reading this article. We are very grateful to Libo Shan (Texas A&M U) for the *A. thaliana serk* mutant seeds, Rob Swanson (Valparaiso U) for the Col-0 *Lat52p:GUS* seeds, Alice Cheung (UMass Amherst) for the *A. thaliana fer* seeds, Scott Woody (UW Madison) for the *B. rapa* FPsc seeds, Stephen Wright (U Toronto) for the *C. rubella* seeds, and the Arabidopsis Biological Resource Center (ABRC) for the *E. salsugineum* seeds. We are also thank Moritz Nowack (Ghent U) for the stigmatic papillae RNA-Seq dataset, Christopher Grefen (Ruhr-U Bochum) for the 2in1 BiFC vectors, Darrell Desveaux (U Toronto) for the Y2H vectors, and the ABRC for the gateway entry vectors with the cytosolic kinase domains. H.K.L was supported by an Ontario Graduate Scholarship (OGS), and this research was supported by a grant from Natural Sciences and Engineering Research Council of Canada to D.R.G.

## Author contribution statement

H.K.L and D.R.G conceived the project, designed the experiments, analyzed the data, and wrote the manuscript. H.K.L. conducted all of the experiments. Both authors commented on the manuscript before submission.

## Materials and Methods

### Plant materials and growth conditions

All seeds for *A. thaliana* (Col-0, transgenic lines, T-DNA insertion mutants and CRISPR deletion mutants), *E. salsugineum, C. rubella*, and *N. benthamiana* were sterilized and cold treated for at least 2 days at 4 °C. Then, they were either transferred directly onto soil or germinated on ½ Murashige and Skoog medium plates with 0.4% (w/v) phytoagar at pH 5.8 at 22°C under 16h light. The 7 days old seedlings were transferred to soil supplemented with 1g/L of Plant Prod All Purpose Fertilizer 20-20-20 and placed in growth chambers under a 16-h-light/8-h-dark cycle at 22°C. All the mutants used in this study are listed in Supplementary Table 6 (CRISPR-generated mutants and SALK identifiers for the T-DNA mutants [35, 38]). See Supplementary Table 5 for the primers used to genotype the *RK* mutants.

### Plasmid construction and plant transformation

The *A. thaliana RK* gene list comprising of 472 genes was retrieved [52, 53], and expression profiles were examined in the TRAVA developmental transcriptome RNA-Seq dataset [31], a stigmatic papillae RNA-seq dataset [54], and two stigma microarray datasets [55]. Based on their expression profiles, 55 RKs were chosen for the Y2H screen (Supplementary Table S1). Entry vectors with cytosolic kinase domains for 40 LRR-RKs were generated by Gou *et al* [56]. The rest of the entry vectors were cloned from *A. thaliana* Col-0 ecotype cDNA using pCR™8/GW/TOPO® TA Cloning® Kit (ThermoFisher Scientific) or by using Gateway® BP Clonase™ II Enzyme Mix (ThermoFisher Scientific) and pDONR207 (Addgene). At-BKN1 was cloned from Hh-0 ecotype and other Brassicaceae RK and BKN cDNAs were cloned from *B. rapa* FPsc, *E. salsugineum* (Thellungiella halophila) [57] and *A. lyrata* ssp. *petrea* [58]. All of the entry vectors created were gateway cloned into their appropriate destination vectors (pEG202, pJG4-5 or pBiFCt-2in1-CC) by using Gateway™ LR Clonase™ II Enzyme Mix (ThermoFisher Scientific). CRISPR vectors carrying two guide RNAs were cloned into pBEE401E as previously described [29, 30, 59] and transformed into *A. thaliana* via *Agrobacterium*-mediated floral dip method [60]. T1 seeds were germinated on soil and selected from spraying Basta herbicide, which was then followed with PCR screens to detect for the herbicide marker, deletions, and wild-type genes.

### Duplex-A Yeast Two-Hybrid assay

The Y2H assay was conducted using the DupLEX-A yeast two-hybrid system (OriGene). The EGY48 yeast strain was transformed with pJG4-5 carrying RK kinase domains or BKNs (prey) and the RFY206 yeast strain was transformed with pEG202 carrying BKNs (bait) or RK kinase domains with pSH18-34 (LacZ reporter plasmid) as described in [61]. The two strains were mated on YPD plates, transferred onto YNB-UHT selection plates to allow for growth and selection. Diploid colonies were then transferred onto either YNB(galactose)-UHT+X-gal or YNB(glucose)-UHT+X-gal and grown for 48∼72 hours period to assess β-galactosidase activity.

### BiFC and confocal microscopy

The 2in1-bimolecular fluorescence complementation assay system developed by Grefen and Blatt[50] was used. The full length RK and BKN cDNAs were cloned into the pBiFCt-2in1-CC gateway vector. *Agrobacterium* GV2260 was transformed with the BiFC vectors by electroporation and then used to infiltrate leaves of 5 weeks old *N. benthamiana* [62]. CLSM images of abaxial leaf epidermis of *N. benthamiana* were taken 48-hours after *Agrobacterium*-mediated infiltration. Images were taken using Leica TCS SP5 confocal microscope with the following excitation intensities and bandpass settings: YFP: 514nm and 521-553nm. RFP: 543nm and 560-615nm. Leica LAS AF lite software was used for further image processing.

### Pollen hydration, aniline blue stain, blue dot assays and silique clearing

To manually pollinate mutant stigmas or Col-0 stigmas with Col-0 or Col-0 *Lat52p:GUS* pollen grains, stage 12 flower buds [63] were emasculated and wrapped with plastic wrap. All pollinations were conducted under humidity levels that were lower than 50% for more consistent results.

#### Pollen hydration assay

At 24-hours post-emasculation, the pistils with fully elongated papillae were mounted on ½ Muraskige and Skoog plate. A single anther from Col-0 were used to lightly hand-pollinate the stigma and pictures were taken at 0- and 10-minutes post-pollination by using Nikon sMz800 microscope with 1.5x objective at 6x magnification. Ten random pollen grains adhered to the stigma were selected and diameters of pollen grains were measured at 0- and 10-minutes using NIS-elements imaging software. Three pistils were used (n=30) per mutant.

#### Aniline blue stain

At 24-hours post-emasculation, pistils with fully elongated papillae were lightly pollinate using a single Col-0 anther per pistil. At 2-hours post-pollination, the pistils were submerged in 300μL fixative (3:1 ethanol: glacial acetic acid) at room temperature for at least 30min. After incubation, pistils were then submerged in 0.5M NaOH at 60°C for 1hr, followed by a series of three washes with 500μL of sterile water. Finally, pistils were stained with 500μL of 0.1% (w/v) aniline blue at room temperature and was mounted on a slide with Vectashield (Vector Laboratories, H-1400) to prevent photobleaching. Brightfield and aniline blue images were taken using Zeiss Axioskop2Plus fluorescence microscope.

#### Interspecies pollinations

Previously reported experimental procedures [9] were followed with the pollinated samples left for 6 hours prior to fixing and aniline-blue staining. Similarly, compatibility scores were calculated as previously described [9] where the numbers of pollen tubes in the styles of the aniline-blue images were counted and assigned one of the following scores: 1, no tubes; 2, 1-19 tubes; 3, 20-39 tubes; 4, 40-59 tubes; 5, ≥60 tubes.

#### Blue dot assay

At 24-hours post-emasculation, pistils with fully elongated papillae were lightly pollinates using a single Col-0 *Lat52p:GUS* [44] anther per pistil. At 2- and 6-hours post-pollination pistils were submerged in 500μL fixative (80% acetone) at room temperature overnight. Pistils were then submerged in the X-gluc solution (5mM Potassium Ferrocyanide, 5mM Potassium Ferriccyanide, 50mM NaPO4 pH7, 0.5mg/ml X-gluc (5-bromo-4-chloro-3-indolyl-”-D-glucoronic acid, cyclohexylammonium salt) at 37°C overnight [64]. Finally, pistils were mounted on slides with 50% glycerol, and images were taken using Nikon sMz800 microscope with 1.5x objective at 3x magnification.

#### Silique clearing

At 24-hours post-emasculation, pistils with fully elongated papillae were lightly pollinates using double Col-0 anther per pistil. The siliques were harvested 2-weeks post-pollination and were subsequently submerged in the clearing medium (1:1:0.93, Lactic acid:Glycerol:Phenol) for 1-week. The siliques were mounted on slides and imaged using Nikon sMz800 microscope with 1.0x objective at 1x magnification.

### Quantification and Statistical Analysis

Pollen grain diameter, pollen tube length and seed counts were measured using NIS-element software as indicated in the method section. In each experiment, the sample sizes, repeats (n) and P values were indicated in the figure legends. Statistical significance was calculated using SPSS software; One-way ANOVA with Tukey-HSD post-hoc test.

### Sequence Data Availability

Sequence data from this report can be found in TAIR or GenBank/EMBL databases under the following accession numbers: RKF1, At1g29750; RKFL1, At At1g29720; RKFL2, At1g29730; RKFL3, At1g29740; LIK1, At3g14840; BSR440, At1g53440; BSR430, At1g53440; BSR650, At1g07650; At-BKN, At5g11400; At-BKN2, At5g11410, Al-lyr-BKN1, AL6G22040; At-CST, At4g35600; SERK1, At1g71830; SERK2, At1g34210; SERK3, At4g33430; SERK4, At2g13790; SERK5, AT2g13800; Br-RKF1 A8, XM_009111396.2; Br-RKFL1 A8, XM_009111402.2; Br-RKFL1 A9, XM_009116971.2; Br-RKFL1 A2, XM_009126838.2; Es-RKF1, Thhalv10006670m; Al-RKF1, AL1G43580.t1

## Supplemental Information

**Figure S1.**
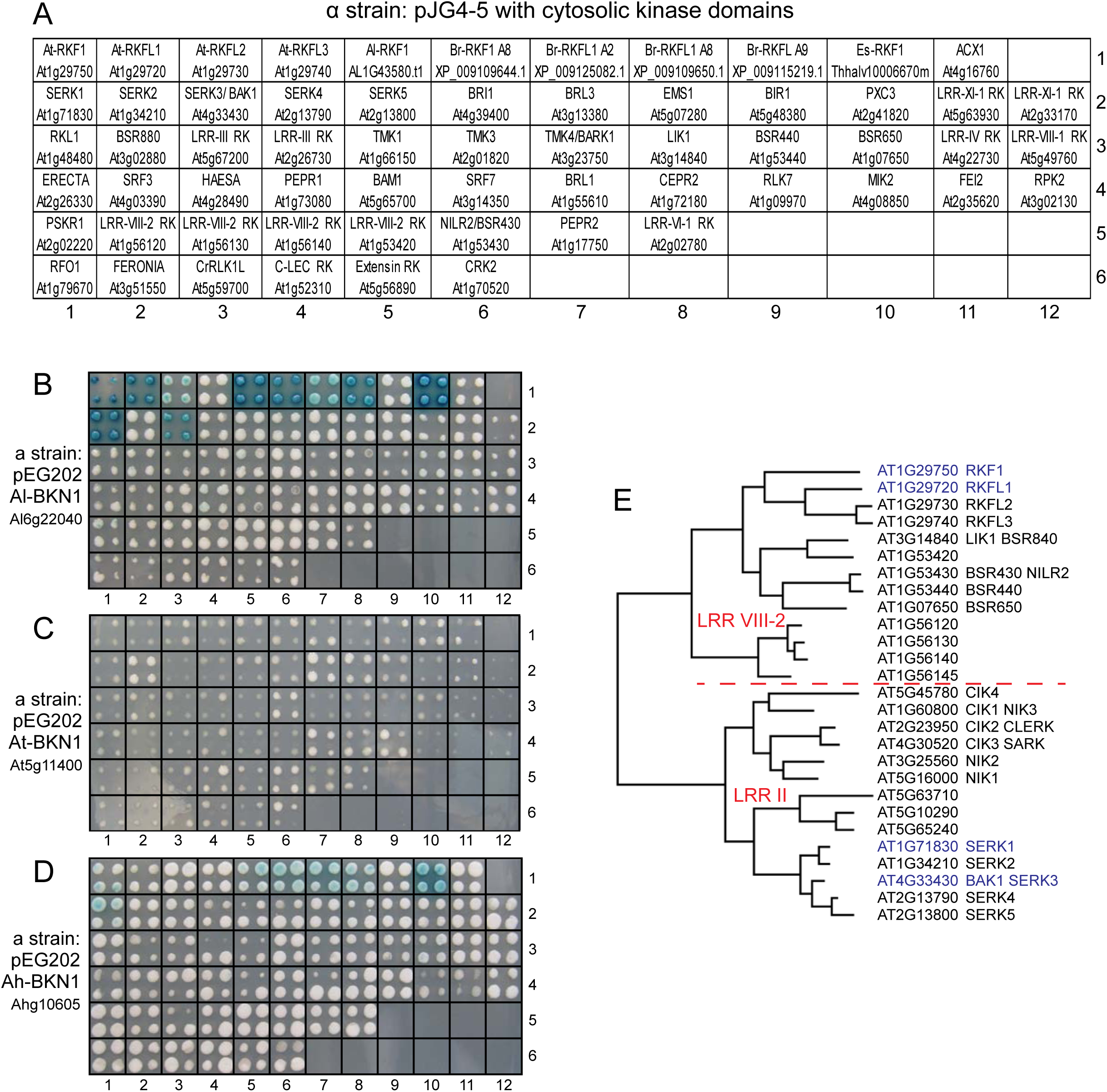
Pairwise Y2H interaction screen of the RK cytosolic kinase domain library with different BKN1 proteins. (A) Shows the key with all the RK gene names. The EGY48 yeast strain was transformed with the pJG4-5 prey vector containing cytosolic kinase domains from different plant RKs to generate the RK cytosolic kinase domain library. ACX1 was also included as a negative control. These transformants were then mated with the RFY206 yeast strain carrying the pEG202 bait vector with the indicated BKN protein: (B*) A. lyrata ssp. petrea* BKN1 (Al-BKN1), (C) *A. thaliana* Hh-0 BKN1 (At-BKN1) and (D) *A. halleri* BKN1 (Ah-BKN1). Mated yeast colonies were pinned in quadruplicate on YNB-UHT + galactose & X-gal media and observed for *LacZ* reporter activity after 48-hrs (80-hrs for Ah-BKN1). (E) Phylogeny of the LRR VIII-2 and LRR II RKs that contained interacting RKs with Al-BKN1 and Ah-BKN1. The phylogeny of these 2 clades was generated in Mega-X (Kumar et al. 2018. Mol Biol Evol 35:1547-1549) and is based on the LRR-RLK maximum likelihood phylogeny published by Mott et al. 2016. Genome Biol.17:98.

**Figure S2.**
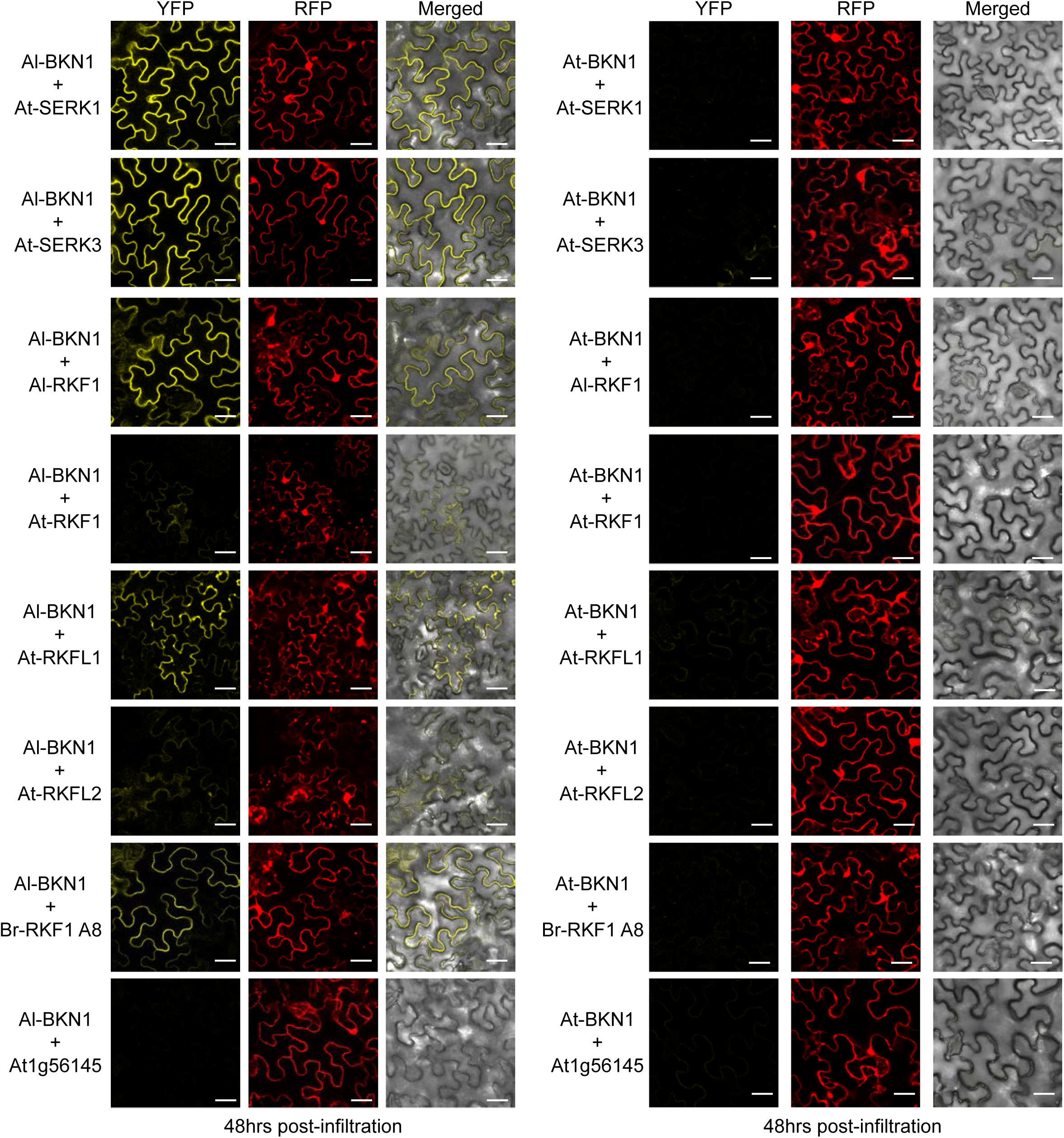
Pairwise interaction assay testing Brassicaceae RKF1 homologues and SERKs with different BKN1 proteins in the BiFC assay. Full length RKF1 homologues and BKN1s were cloned with C-terminal fusions of the EYFP halves. CLSM images of abaxial leaf epidermis of *N. benthamiana* were taken 48-hours after *Agrobacterium*-mediated infiltration. The single BiFC vector with the receptor kinase and BKN clones also carries an RFP marker which was used to identify epidermal cells transformed with the BiFC vector. The presence of a YFP signal in the left panel for each set indicates a positive protein interaction. The right panel for each set represents a merged images of the brightfield and YFP images. At1g56145 is another LRR VIII-2 RK that was used as a negative control. Images were taken using the Leica TCS SP5 confocal microscope with the following excitation intensities and bandpass settings: YFP: 514nm and 521-533nm. RFP: 543nm and 560-615nm, Grefen and Blatt. 2012. Biotechniques.311-314; Scale bar = 30 µm.

**Figure S3.**
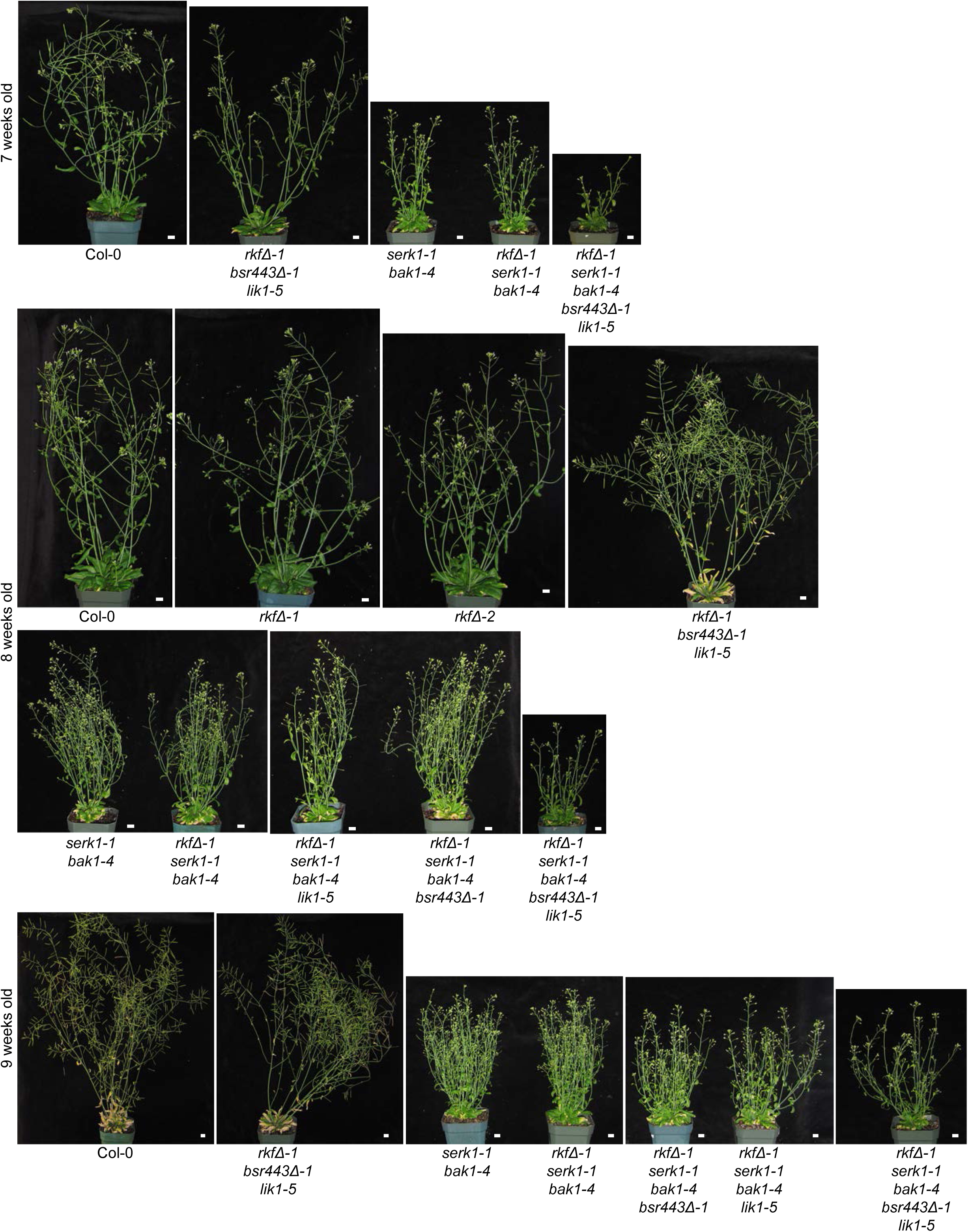
Representative images of flowering *A. thaliana* wild-type Col-0 and *RK* mutant plants. Whole plant photos were taken at 7, 8 and 9 weeks and show the delayed flowering in mutant carrying the *serk-1bak1-4* mutations. Scale bar = 1.0 cm.

**Figure S4.**
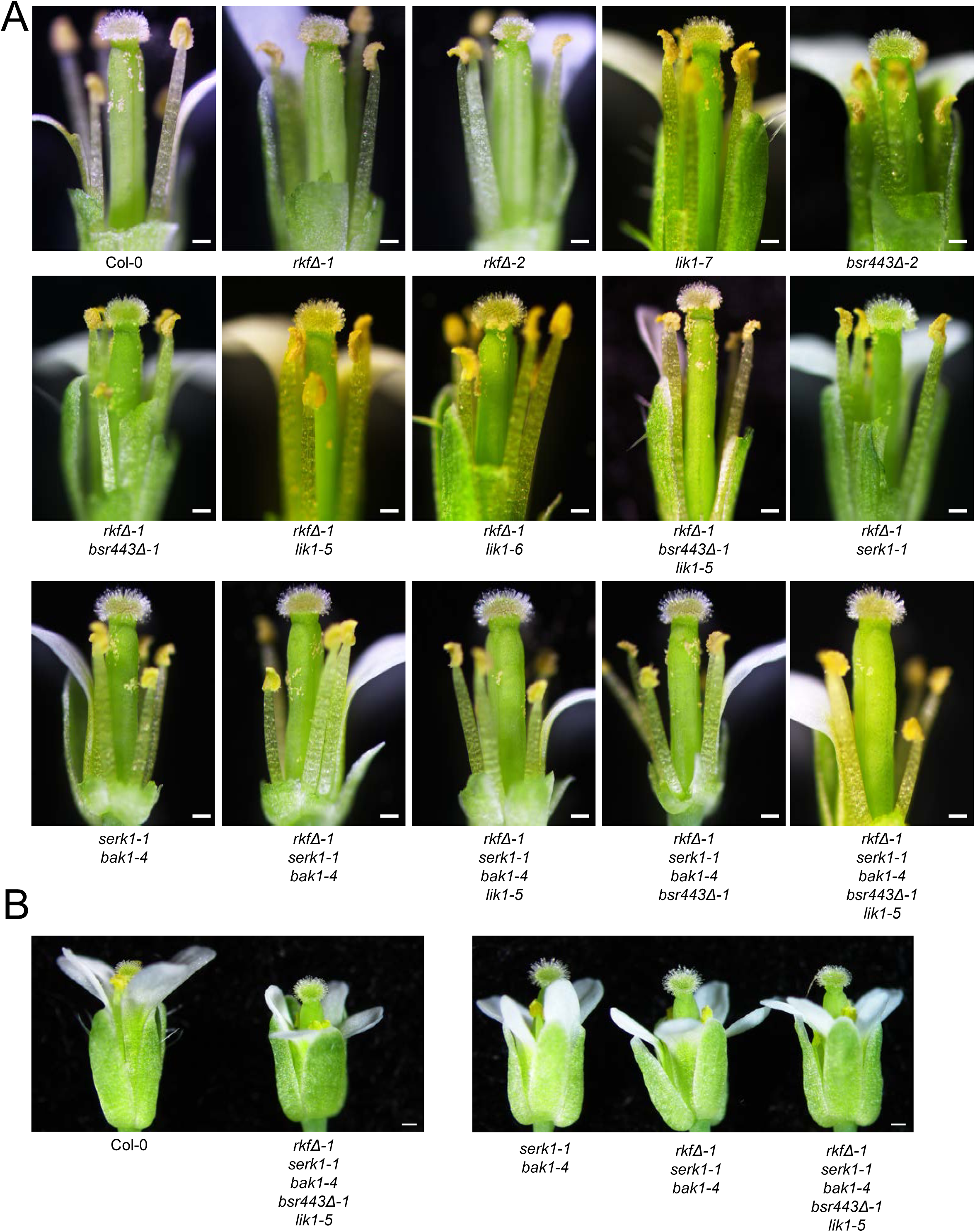
Representative images of stage-14 flowers from *A. thaliana* wild-type Col-0 and *RK* mutant plants. (A) From A. thaliana wild-type Col-0 and the different *RK* combination mutant plants (some petals and sepals have been removed to better show the pistil and anthers). (B) Select flowers showing the full flower view to show that the pistils are similar in size. Scale bar = 200 µm

**Figure S5.**
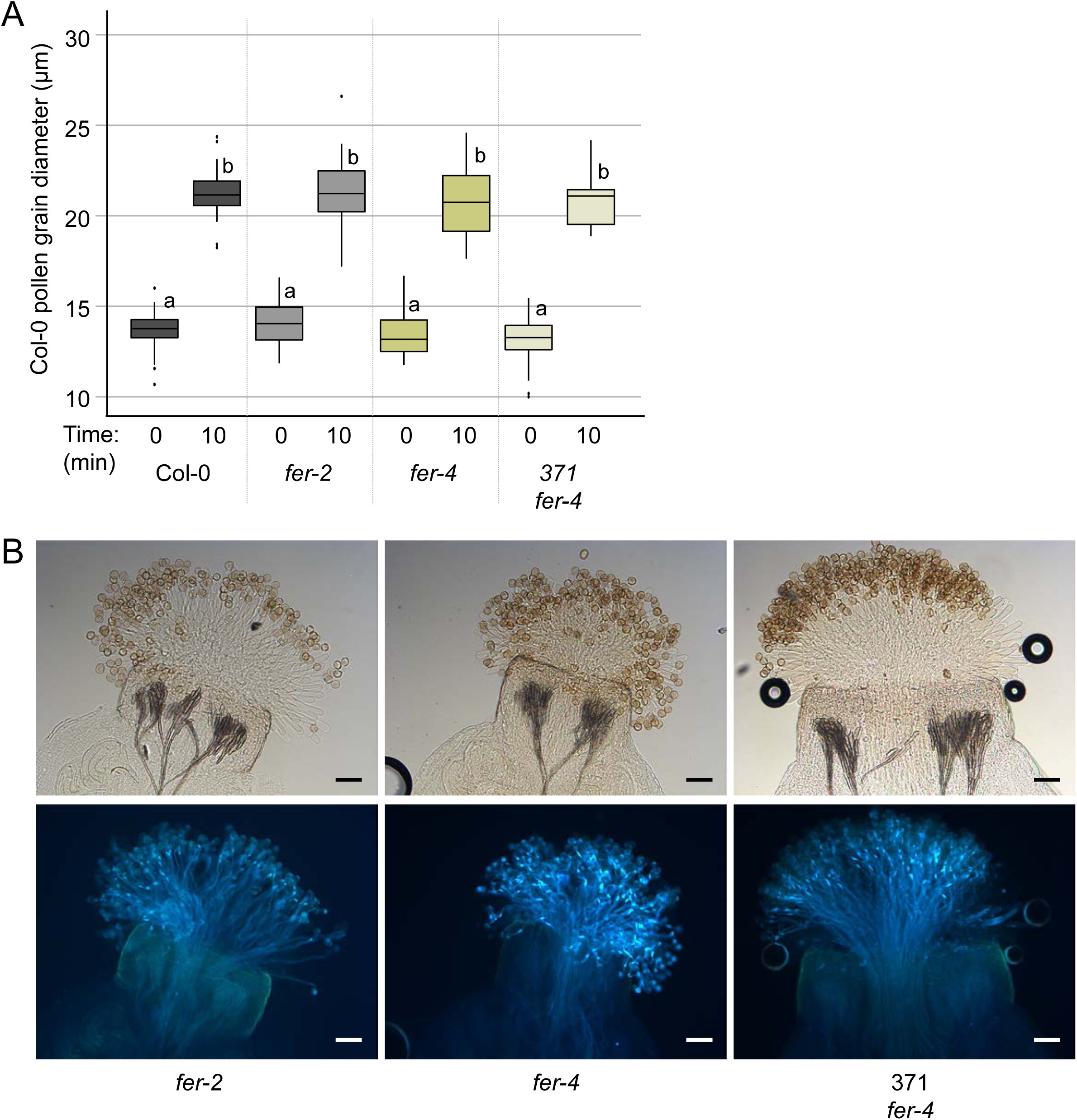
Pollen hydration and aniline blue assay for the *fer* mutant. (A) Col-0 pollen hydration assay. Col-0 pollen grain diameters were measured at 0-and 10-minutes post-pollinations on stigmas from Col-0, *fer-2, fer-4* and the rescued *fer* line (371 *fer-4*). Data are plotted as box plots displaying first (Q1) and third (Q3) quartiles split by the median; the whiskers extend maximum of 1.5 times the interquartile range beyond the box. Outliers are indicated as black dots. n=30 pollen grains per line, P<0.05 (One-way ANOVA with Tukey-HSD post-hoc test). (B) Aniline blue stained pistils 2-hrs post-pollination. Col-0 pollen was applied to pistils from Col-0, *fer* mutants and the rescued *fer* line (371 *fer-4*). Brightfield images on top and aniline blue images on bottom. Scale bar= 100 µm.

**Figure S6.**
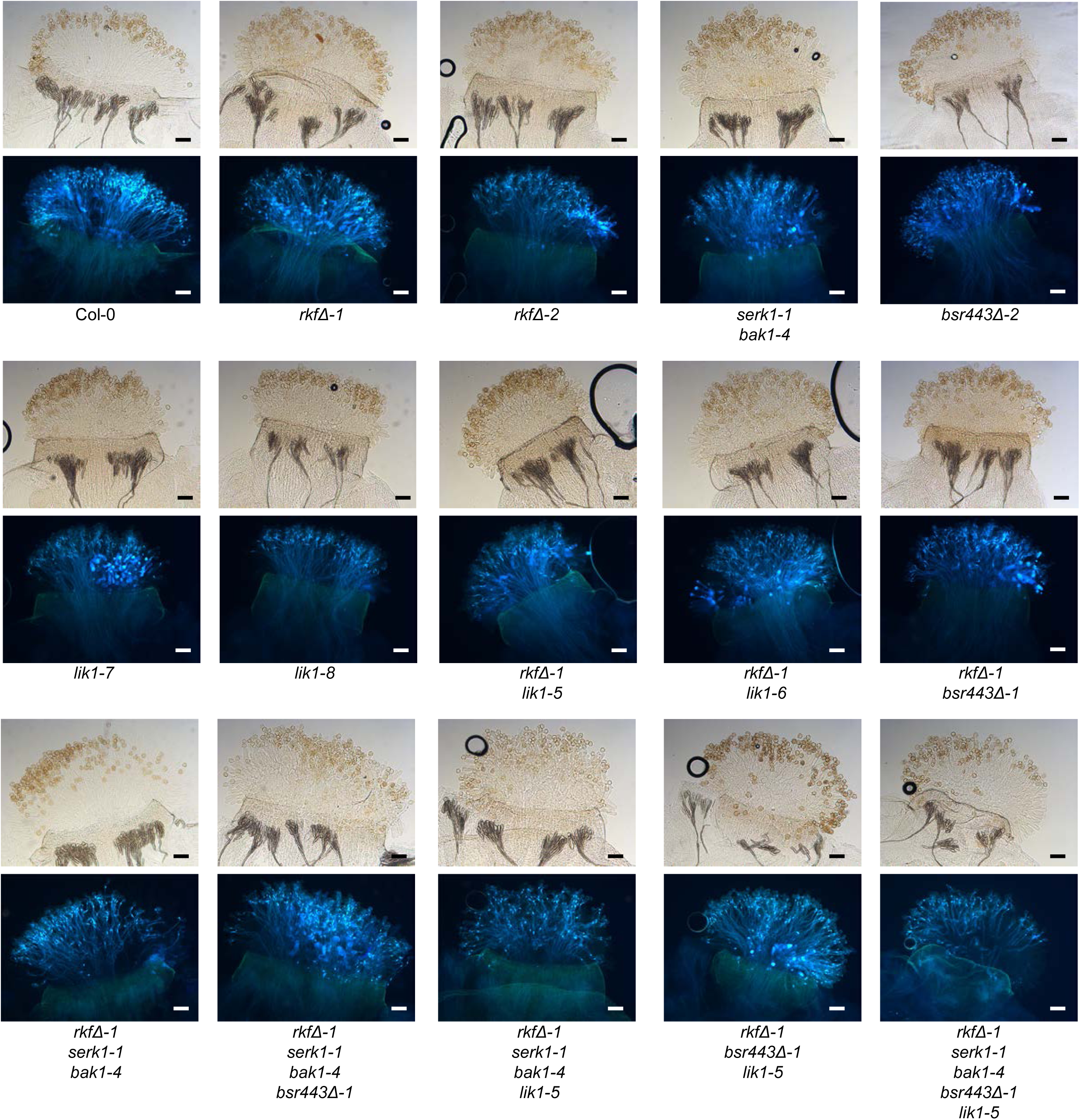
Representative images aniline blue stained pistils from *A. thaliana* wild-type Col-0 and *RK* mutant plants pollinated with Col-0 pollen. Col-0 pollen was applied to stigmas from wild-type Col-0 and the different *RK* mutant pistils. Pistils were collected at 2-hours post-pollinations and fixed and stained with aniline blue. All plants were homozygous mutants for the indicated mutant alleles, and for each genotype, brightfield images are shown on the top and aniline blue stained images are shown on the bottom. Scale bar= 100 µm.

**Figure S7.**
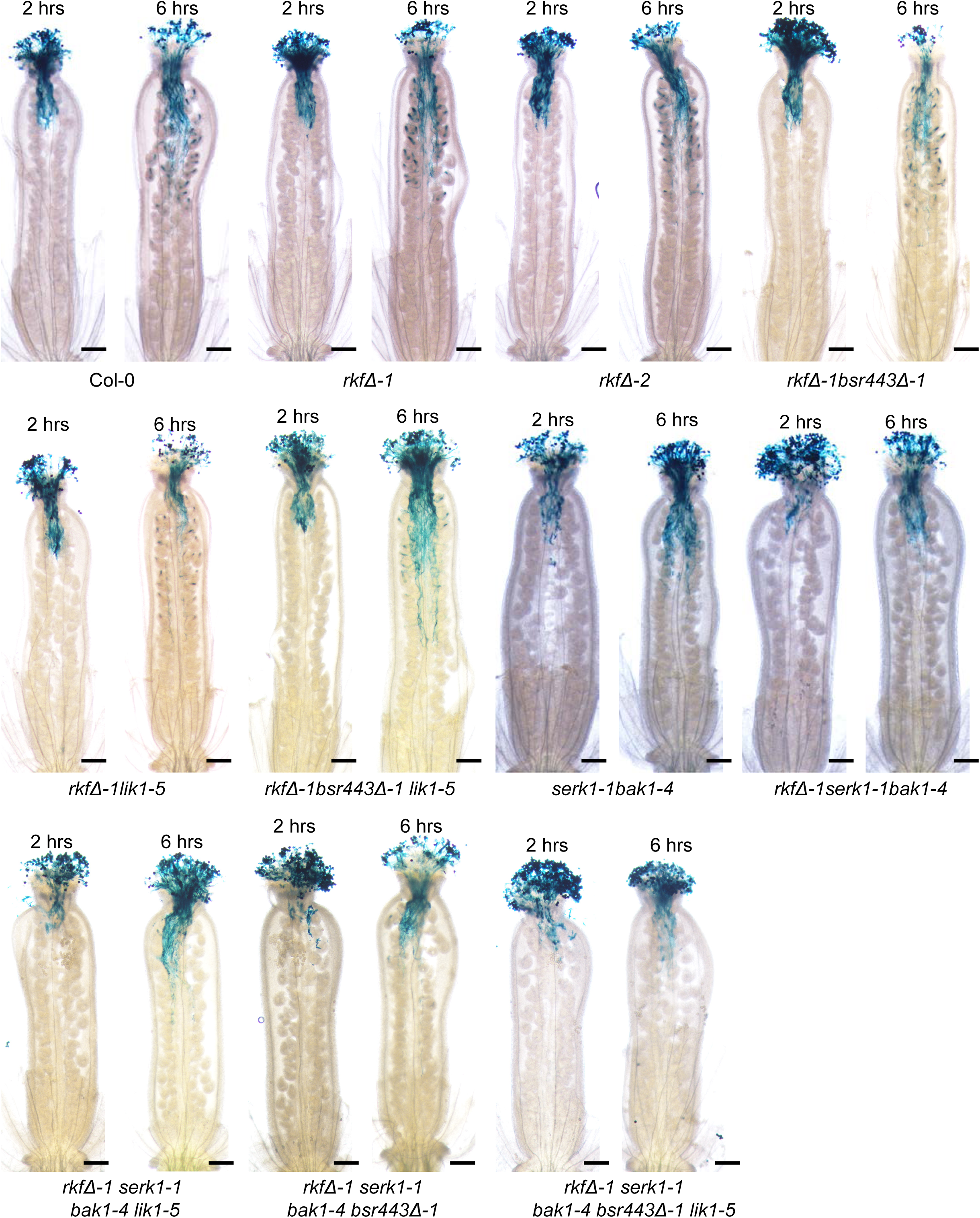
Representative images of Col-0 and *RK* mutant pistils pollinated with Col-0 *Lat52p::GUS* pollen. Col-0 *Lat52p::GUS* pollen was applied to pistils from Col-0 or *RK* mutants and were fixed at either 2- or 6-hr post-pollinations. Scale bar = 200 µm.

**Table S1. eNorthern with the Klepikova RNA-Seq for 468 Arabidopsis RK genes.**

**Table S2.** Summary of Expression Patterns for the Selected Receptor Kinases from Different Expression Datasets.

|  | Gene Name | Extracellular Domain | AGI | Stigmatic Papillae |  | Stigmas |  |  |  | TRAVA Stigma | GUS activity |  |  |
| --- | --- | --- | --- | --- | --- | --- | --- | --- | --- | --- | --- | --- | --- |
|  |  |  |  | T1 Av | T1 SE | 0 min Pol Average | Coat SE | 15 min Pol Average | Coat SE |  | Stigma | Style |  |
| 1 | FERONIA | CrRLK1L | At3g51550 | 667 | 14 | 2005 | 32 | 2181 | 137 | 7036 | Yes* | Yes* | * Duan et al. 2010 |
| 2 | BRI1 | LRR-Xb-1 | At4g39400 | 617 | 37 | 842 | 97 | 796 | 95 | 4925 | Yes | Yes |  |
| 3 | PXC3 | LRR-XI-1 | At2g41820 | 553 | 19 | 292 | 47 | 237 | 89 | 9778 | Yes | Yes |  |
| 4 | RKL1 | LRR-III | At1g48480 | 483 | 44 | 2366 | 411 | 1911 | 550 | 3649 | No | Weak |  |
| 5 | BRL3 | LRR-Xb-1 | At3g13380 | 327 | 7 | 540 | 136 | 528 | 110 | 1632 | Yes | Yes |  |
| 6 | TMK1 | LRR-IX | At1g66150 | 276 | 2 | 1380 | 158 | 1282 | 207 | 3379 | Yes | Yes |  |
| 7 |  | LRR-IV | At4g22730 | 211 | 4 | 463 | 109 | 431 | 100 | 2638 | Yes | Yes |  |
| 8 |  | LRR-VIII-1 | At5g49760 | 192 | 10 | 193 | 54 | 187 | 35 | 4006 | Yes | Yes |  |
| 9 | CRK2 | Cysteine-rich | At1g70520 | 175 | 4 | 830 | 268 | 937 | 200 | 725 | - | - |  |
| 10 |  | LRR-VI-1 | At2g02780 | 160 | 10 | 418 | 45 | 428 | 54 | 2655 | Yes | Yes |  |
| 11 | LIK1/BSR840 | LRR-VIII-2 | At3g14840 | 150 | 15 | 180 | 41 | 245 | 21 | 4451 | Yes | Yes |  |
| 12 | TMK3 | LRR-IX | At2g01820 | 147 | 8 | 1183 | 182 | 1069 | 97 | 2104 | Yes | Yes |  |
| 13 |  | LRR-XI-1 | At5g63930 | 141 | 1 | 222 | 8 | 238 | 30 | 577 | No | Weak |  |
| 14 | TMK4/BARK1 | LRR-IX | At3g23750 | 141 | 6 | 517 | 108 | 464 | 49 | 2085 | Yes | Yes |  |
| 15 | EMS1 | LRR-Xb-1 | At5g07280 | 133 | 8 | 222 | 25 | 184 | 22 | 1903 | Weak | Yes |  |
| 16 |  | LRR-XI-1 | At2g33170 | 132 | 8 | 323 | 71 | 280 | 19 | 1288 | Weak | Yes |  |
| 17 | BIR1 | LRR-Xa | At5g48380 | 114 | 4 | 300 | 41 | 452 | 33 | 1721 | No | Weak |  |
| 18 | BSR440 | LRR-VIII-2 | At1g53440 | 108 | 2 | 248 | 46 | 332 | 28 | 1687 | Yes | Yes |  |
| 19 | ERECTA | LRR-XIIIb | At2g26330 | 103 | 4 | 543 | 123 | 503 | 86 | 1704 | No | Weak |  |
| 20 | BSR880 | LRR-III | At3g02880 | 101 | 2 | 276 | 64 | 374 | 54 | 974 | No | Weak |  |
| 21 | SRF3 | LRR-V | At4g03390 | 96 | 2 | 329 | 72 | 265 | 42 | 1159 | Yes | Yes |  |
| 22 |  | Extensin | At5g56890 | 93 | 7 | 192 | 55 | 147 | 12 | 1928 | - | - |  |
| 23 | HAESA | LRR-XI-1 | At4g28490 | 91 | 5 | 322 | 107 | 335 | 68 | 1063 | Yes | Yes |  |
| 24 | ANJ | CrRLK1L | At5g59700 | 84 | 1 | 226 | 85 | 244 | 92 | 1380 | - | - |  |
| 25 | SERK3/BAK1 | LRR-II | At4g33430 | 73 | 2 | 359 | 51 | 419 | 57 | 1417 | Yes | Yes |  |
| 26 | PEPR1 | LRR-XI-1 | At1g73080 | 72 | 12 | 1641 | 202 | 2581 | 428 | 2052 | Yes | Yes |  |
| 27 | BAM1 | LRR-XI-1 | At5g65700 | 70 | 4 | 294 | 71 | 316 | 38 | 2560 | Weak | Yes |  |
| 28 | SRF7 | LRR-V | At3g14350 | 62 | 5 | 83 | 6 | 100 | 12 | 1175 | Weak | Yes |  |
| 29 | BRL1 | LRR-Xb-1 | At1g55610 | 54 | 2 | 96 | 15 | 94 | 16 | 931 | Yes | Yes |  |
| 30 |  | LRR-III | At5g67200 | 53 | 2 | 91 | 5 | 103 | 23 | 798 | Yes | Yes |  |
| 31 |  | LRR-III | At2g26730 | 52 | 2 | 74 | 1 | 74 | 4 | 1491 | No | Weak |  |
| 32 | CEPR2 | LRR-XI-1 | At1g72180 | 50 | 3 | 1288 | 144 | 1154 | 155 | 705 | Yes | Yes |  |
| 33 |  | C-LEC | At1g52310 | 50 | 1 | 329 | 39 | 305 | 35 | 1872 | - | - |  |
| 34 | RFO1 | WAKL22 | At1g79670 | 40 | 2 | 81 | 4 | 138 | 19 | 442 | - | - |  |
| 35 | RKF1 | LRR-VIII-2 | At1g29750 | 39 | 0 | 128 | 12 | 144 | 20 | 1056 | Yes | Yes |  |
| 36 | RLK7 | LRR-XI-1 | At1g09970 | 37 | 7 | 3441 | 290 | 3654 | 366 | 1094 | Yes | Yes |  |
| 37 | MIK2 | LRR-XI-1 | At4g08850 | 35 | 2 | 123 | 16 | 158 | 9 | 863 | Yes | Yes |  |
| 38 | FEI2 | LRR-XIIIa | At2g35620 | 35 | 1 | 66 | 4 | 72 | 7 | 1008 | Yes | Yes |  |
| 39 | RPK2 | LRR-XV | At3g02130 | 34 | 3 | 124 | 24 | 192 | 17 | 715 | No | Weak |  |
| 40 | PSKR1 | LRR-Xb-1 | At2g02220 | 30 | 1 | 64 | 13 | 81 | 10 | 452 | Yes | Yes |  |
| 41 |  | LRR-VIII-2 | At1g56140 | 30 | 3 | 111 | 18 | 209 | 16 | 249 | No | No |  |
| 42 | SERK1 | LRR-II | At1g71830 | 24 | 0 | 271 | 22 | 245 | 9 | 639 | Weak | Yes |  |
| 43 |  | LRR-VIII-2 | At1G56145 | 15 | 0 | 62 | 3 | 69 | 6 | 359 | No | No |  |
| 44 | NILR2/BSR430 | LRR-VIII-2 | At1g53430 | 13 | 1 | 60 | 2 | 101 | 14 | 562 | Yes | Yes |  |
| 45 | SERK2 | LRR-II | At1g34210 | 11 | 1 | 447 | 29 | 417 | 50 | 878 | Yes | Yes |  |
| 46 | SERK5 | LRR-II | At2g13800 | 7 | 0 | 71 | 4 | 122 | 16 | 120 | No | No |  |
| 47 | BSR650 | LRR-VIII-2 | At1g07650 | 7 | 1 | 124 | 29 | 149 | 26 | 718 | Yes | Yes |  |
| 48 | RKFL1 | LRR-VIII-2 | At1g29720 | 6 | 1 | 38 | 1 | 37 | 1 | 697 | No | Weak |  |
| 49 | SERK4 | LRR-II | At2g13790 | 5 | 0 | 72 | 8 | 146 | 11 | 147 | No | Weak |  |
| 50 | RKFL2 | LRR-VIII-2 | At1g29730 | 4 | 1 | 42 | 1 | 39 | 1 | 62 | No | Weak |  |
| 51 |  | LRR-VIII-2 | At1g56120 | 2 | 0 | 25 | 2 | 26 | 2 | 56 | No | No |  |
| 52 |  | LRR-VIII-2 | At1g56130 | 1 | 0 | 30 | 1 | 27 | 1 | 59 | Weak | Yes |  |
| 53 | RKFL3 | LRR-VIII-2 | At1g29740 | - | - | 35 | 1 | 38 | 2 | 25 | No | No |  |
| 54 |  | LRR-VIII-2 | At1g53420 | - | - | 22 | 2 | 21 | 1 | 0 | Weak | Weak |  |
| 55 | PEPR2 | LRR-XI-1 | At1g17750 | - | - | 63 | 8 | 168 | 33 | 54 | Yes | Yes |  |

**Table S3.**
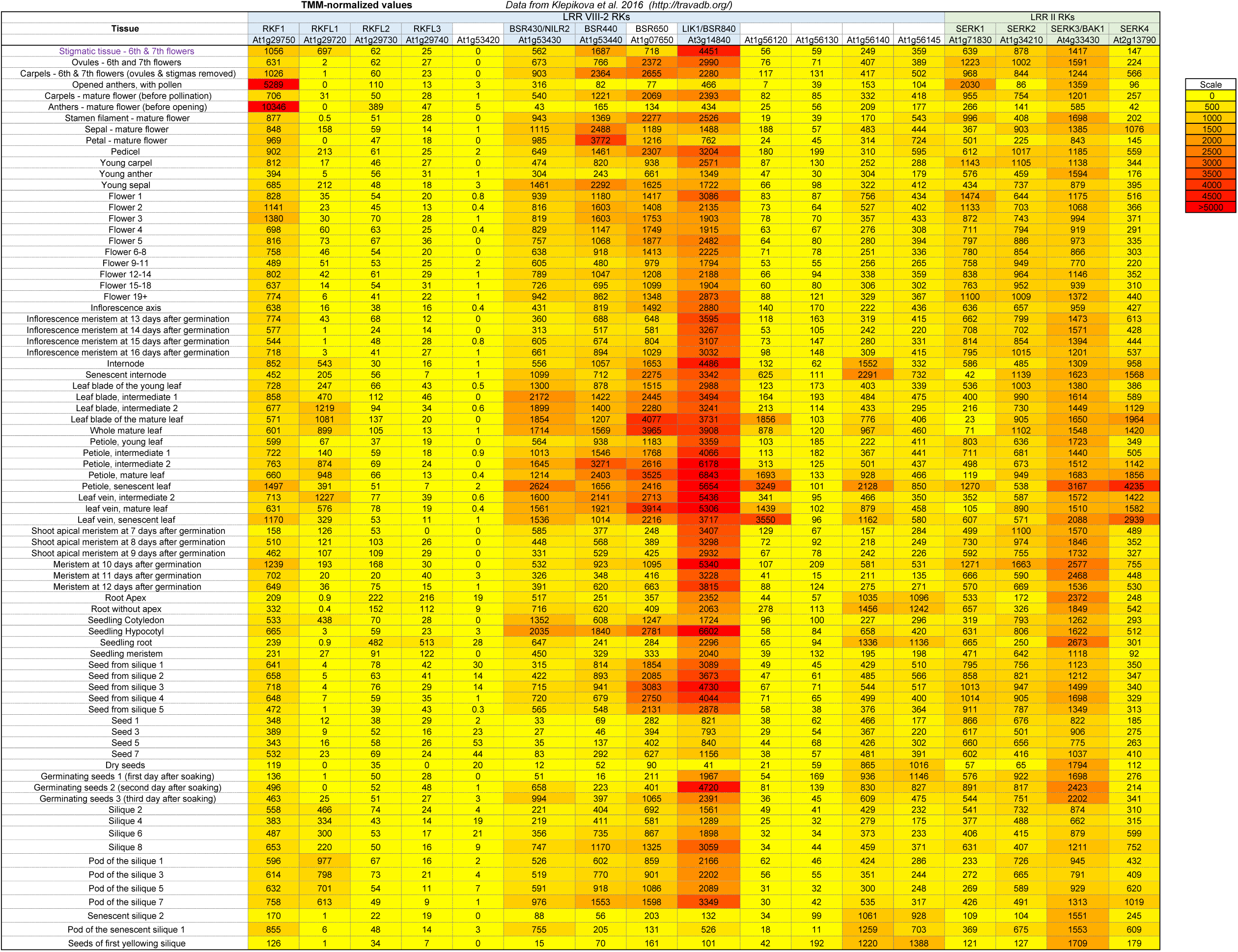
eNorthern with the Klepikova RNA-Seq for the Arabidopsis LRR VIII-2 RK genes and the LRR II RK genes.

**Table S4. Gateway entry cDNA clones used.**

**Table S5. List of primers used to detect CRISPR or T-DNA deletions 5’->3’.**

**Table S6. T-DNA and CRISPR knockout mutants**

